# Neural evidence for action-related somatosensory predictions

**DOI:** 10.1101/2024.11.29.626056

**Authors:** Caoimhe Moran, Hinze Hogendoorn, Ayelet N. Landau

## Abstract

The tactile consequences of self-initiated movements are thought to be predicted by a forward model, yet the precise neural implementation of these predictions remains unclear. In non-motor contexts, expectations are thought to activate sensory neurons tuned towards the expected stimulus. This acts as a predictive template against which afferent sensory input is compared. It is unclear whether forward model predictions have a similar neural instantiation. Here we employed time-resolved multivariate decoding on human electroencephalography (EEG) during self-generated movements to examine the content of predictive neural activity. Human participants performed index finger movements which were predictably paired with a vibration to either the index or ring finger of the opposite, passive hand. On some trials the tactile stimulus was unexpectedly omitted. Results revealed above-chance finger decoding in the pre-movement period supporting a predictive representation of expected stimulation location. As the movement approached, this predictive activity became similar to late-stage processing of a physical tactile stimulus. On omission trials, we found that despite the absence of afferent input, finger location could be decoded ∼120 ms after expected stimulus onset. This shows a stimulus-specific omission response. Together these findings indicate that self-generated movement pre-activates neurons tuned towards expected tactile consequences.

## 1 Introduction

Every movement comes with an expectation of its sensory consequences. For example, when you kick a ball, various different sensations are anticipated such as the feeling of the ball’s weight against your foot, the sight of the ball moving through the air and the sense of your leg as it moves through space. These expectations act as templates against which incoming sensory information can be compared, facilitating optimal engagement with the environment (de Lange et al., 2018; Summerfield & Egner, 2009). This idea dates back to William James who argued that all voluntary actions are preceded by a prediction of their consequences, termed action-effect predictions (James, 1890).

Major theories on action-effect prediction propose that the brain uses a forward model to predict the sensory consequences of self-produced actions (Blakemore et al., 1999; Dogge et al., 2019; Kilteni & Ehrsson, 2020). As a motor command is sent to motor control areas, a copy of that command, known as corollary discharge, is sent to the forward model; a system capable of simulating body movements in order to anticipate the resulting sensory consequences (Todorov, 2004). It is thought that incoming sensory signals are continuously compared to sensory predictions allowing a Bayes-optimal estimate of action-effects. Errors made when predicting afferent inputs are exploited by the system to update internal models, refining future predictions. By their nature, self-generated actions have highly predictable consequences resulting in little to no prediction error (PE). This predictive mechanism explains extensive evidence of sensory attenuation for self-generated action consequences across multiple domains, including vision (Hughes & Waszak, 2011; Straube et al., 2017), audition (Baess et al., 2009; Nelson et al., 2013; Schneider et al., 2018; Weiss et al., 2011) and touch (Blakemore et al., 1999; Kilteni & Ehrsson, 2020). For example, self-produced touch is often perceived as less intense or ticklish (Weiskrantz et al., 1971; Blakemore et al., 1999, 2000; Shergill et al., 2003; Bays et al., 2005, 2006; Kilteni et al., 2019; Job & Kilteni, 2023) and results in reduced activation in the somatosensory cortex and cerebellum (Blakemore et al., 1999; Hesse et al., 2010; Shergill et al., 2013; Kilteni & Ehrsson, 2020, 2020, 2023). These findings indicate that predictive mechanisms play a role in modulating the response to self-generated sensations.

On the flip side of attenuated neural responding for fulfilled predictions, prediction violations lead to an enhanced neural response. The mismatch negativity (MMN) is a component of the event-related potential (ERP) that captures detection of prediction violations (Alho, 1995) and is widely regarded as the neural correlate of prediction error, primarily studied in the auditory domain (Waszak & Herwig, 2007; Korka et al., 2019). Both neural attenuation and the MMN are used as key evidence in support of predictive coding (Clark, 2013). However, alternative explanations that do not rely on predictive mechanisms cannot be ruled out. For example, neural adaptation – characterised by reduced activity in stimulus-specific neurons due to synaptic depression or lateral inhibition from repeated stimulus exposure - could also account for these findings (May & Tiitinen, 2001; Jääskeläinen et al., 2004; May & Tiitinen, 2010). In addition, neural response modulation by stimulus expectation can provide only indirect evidence for the existence of predictions (Schröger et al., 2015). The challenge lies in disentangling endogenous predictions from bottom-up sensory input. A more direct measure of endogenous predictions could be achieved by removing afferent sensory input entirely (Arnal & Giraud, 2012).

Stimulus omission paradigms may offer a better insight into the predictive architecture underlying actions and their sensory consequences. It has been reported that when an action is paired with an auditory stimulus and the stimulus is unexpectedly omitted, sensory like ERP responses are produced, despite the absence of afferent input (SanMiguel et al., 2013; Dercksen et al., 2020, 2022; Korka et al., 2020). Such omission responses are only elicited when stimulus omissions are rare supporting the notion that they represent the PE (SanMiguel et al., 2013). The absence of a physical stimulus suggests that these neural responses result from internal processes, such as predictive mechanisms. Although omission paradigms are well-established in the auditory domain, they remain relatively unexplored in the somatosensory field. However, a recent study found an EEG omission response when an action-related somatosensory prediction was violated (Dercksen et al., 2024). Specifically, they observed a negative omission-related potential 80-100 ms after the anticipated but absent stimulus, perhaps reflecting a PE signal. Yet, traditional ERP measures provide no information about the content of PE signals. For instance, it is unclear whether the error response contains specific details about the expected stimulus, such as intensity or location, or if it is more general, surprise signal.

An essential aspect of tactile prediction is the precise location of the expected touch. This is crucial in order to distinguish between self-generated and externally-generated sensations. Accurate spatial predictions are also critical for establishing a sense of agency i.e. the feeling of having caused one’s own actions and the effects they produce in the world (Haggard & Tsakiris, 2009). For instance, if you are mindlessly tapping your left arm with your right finger and someone simultaneously taps your shoulder, you can easily distinguish between the self-produced and external taps. This suggests that predictions about self-touch are spatially precise. Early work showed that perturbations introduced to the movement path during self-stimulation lead to higher ratings of ’ticklishness’ (Blakemore et al., 1999). This heightened sensation when the tactile stimulus diverges from its spatially predicted path has been attributed to a spatial PE. While efforts have been made to isolate the specificity of prediction-related neural activity in the auditory domain (Korka et al., 2020; SanMiguel et al., 2013); to our knowledge, capturing an identity-specific somatosensory prediction in the context of action remains unexplored.

In the present study, we used a temporally resolved EEG decoding approach to directly examine the spatial specificity of movement-driven tactile predictions. Participants performed self-timed index finger movements, after which a brief tactile stimulus was delivered to either the index or ring finger of the opposite, passive hand. The site of stimulation remained the same within each block of trials, making it fully predictable within each block, and stimulus onset was determined by participants’ self-timed movements. On some trials the stimulus was omitted. We hypothesised that when participants anticipate a vibration on a specific finger as a result of their movement, the preparation of that movement triggers the predictive activation of a finger-specific tactile representation in the somatosensory cortex. When the expected stimulus is omitted, the discrepancy between the prediction and the bottom-up input, or lack thereof, is propagated back through the cortical hierarchy. Given that there is no bottom up input on omission trials, any above-chance decoding of stimulus location must be a direct consequence of predictive activity, perhaps reflecting the prediction error response.

Furthermore, consistent with anticipatory stimulus templates, formed before expected afferent input, we hypothesise above-chance decoding of the expected finger stimulation location in the pre-stimulus period.

## 2 Materials and Methods

### 2.1 Participants

Data was collected from a total of 39 participants, of which four were excluded due to poor EEG classification performance (less than 52% average decoding accuracy when classifying finger location on passive trials). After exclusion 35 participants remained (23 females; age range = 19-35; mean age = 24; SD = 3 years; 5 left-handed). All observers provided informed consent before beginning the experiment and were compensated with credit points or reimbursed 13 USD per hour. The project was approved by the local ethical committee.

### 2.2 Apparatus and stimuli

The experiment was conducted in MATLAB Version R2021b on a BenQ XL2420Z monitor. The experiment was programmed using the Psychophysics Toolbox extension (Brainard, 1997; Pelli, 1997; Kleiner et al., 2007). A black fixation cross was presented in the middle of the screen at approximately 60 cm from participants’ eyes. Stimuli were delivered using TDK PowerHap Piezo Actuators (9 x 9 x 1.1 mm) connected to a Piezo driver (Boreas – BOS1901 Development kit). The piezo driver appears as a normal audio output to the computer. Thus, tactile stimuli were created by programming an audio sine wave which produced a vibration for 40 ms at 300 Hz. An infrared motion tracker (Leap Motion Controller using the Matleap MATLAB interface) was used to track participants finger movements. When the distal phalange (the tip) of the index finger descended 20 mm from the starting position at trial onset, the target stimulus was delivered. Pilot testing ensured that the stimulus felt like an immediate consequence of the movement. Participants’ hands were visually occluded during the experiment and white noise was continually played through headphones to obscure the sound of the tactile stimulus. A test was run prior to experiment onset to ensure participants could not hear the vibration. The white noise volume was adjusted until the stimulus sound was fully obscured.

### 2.3 Experimental design

Participants were seated in a dimly lit room, their left hand, palm up on the table (see Fig. 1a). Tactors were placed on the apex of the left index and ring fingers. Their right arm lay on an armrest, palm face down, with the hand extended over the edge. The motion tracker was placed directly under the right hand (15 cm from the palm). After set-up, participants received verbal and written task instructions. Depending on block type (passive or active) participants were told to keep their right hand still or move their index finger downward. Under both conditions, participants were told to press a foot pedal with their right foot when they detected a tactile vibration at a lower intensity than the others. In addition, the experimenter demonstrated the correct hand positions and showed the participant how to accurately perform the movement. To ensure accurate calibration between the motion camera and movement, participants performed ten index finger movements and received feedback on whether the movement was recorded or not. Subsequently, participants did a short practice in which their movement was followed by the tactile stimulus.

**Figure 1:**
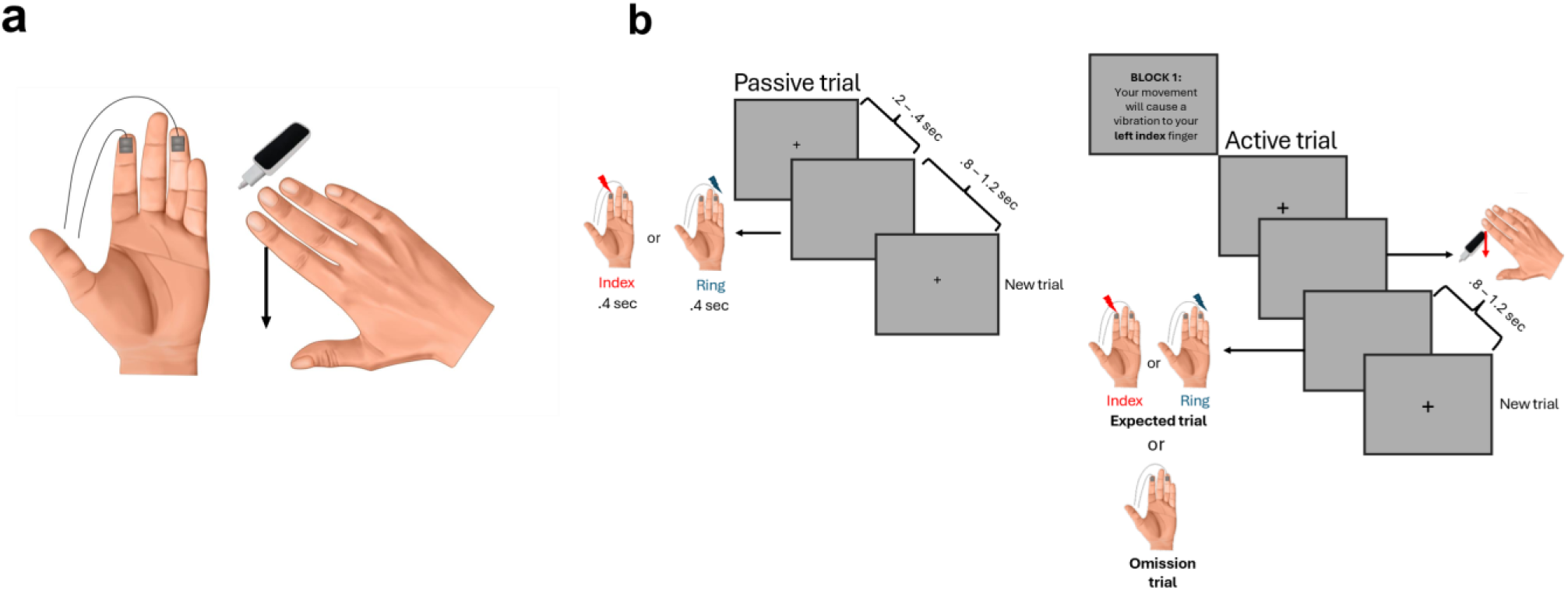
Paradigm and trial types. **(a)** Position of the hands during the experiment. Two tactors were attached to the left index and ring finger. Participants positioned their right hand above the motion sensor camera. The right hand remained still on passive trials while in active trials, participants made a downward movement with the index finger. **(b)** Trial details for passive trials (*left side*) in which participants’ both hands remained still. They received short vibrations (40 ms) to either their left index or ring finger with a jittered inter-trial interval. Active trials (*right side*) required participants to move their right index finger downward. Depending on the block, they received a vibration to their left index or ring finger (expected trials) immediately after the movement. On 20 % of trials the vibration was omitted entirely (omission trials).

Participants completed passive and active blocks. On passive blocks participants looked at the fixation cross at the centre of the screen and remained still. A tactile vibration was delivered to either the left index or ring finger for 40 ms. A jittered interval between 800 and 1200 ms separated each trial and the fixation cross was removed from the screen with each stimulation and presented again at the beginning of the next trial (Fig. 1b).

In active blocks, cued by the appearance of a fixation cross, participants made a downward movement with their right index finger. This was immediately followed by a brief vibration (40 ms) to either the left index or ring finger, termed Active – Expected (AE) trials or omitted entirely, termed Active – Omission (AO) trials. The mapping between the movement and vibration location were counterbalanced across blocks. The stimulus was never delivered to the alternate finger within a block (Fig. 1b).

In total there were 5 passive blocks with 120 trials in each block and 6 active blocks with 80 trials in each block. Omission trials constituted 20% of active trials. Across both active and passive blocks, participants were asked to detect and respond to tactile stimuli at a lower intensity by pressing a foot pedal. These made up 7.5% of trials (6 trials per active block and 10 trials per passive block) and were removed from the main analysis. This task ensured that participants were paying attention to the tactile stimulus but not specifically that it was delivered to the index or the ring finger, which are the conditions that we use for the decoding analysis. All blocks of the same type (passive or active) occurred consecutively. The block order was counterbalanced across participants such that participants completed either the active or passive blocks first.

### 2.4 EEG Acquisition and preprocessing

EEG was recorded throughout the experiment as participants performed the passive and active trials. A g.GAMMAcap (gTec, Austria) and a g.HIamp amplifier (gTec, Austria) were used. The 62 active electrodes on the cap were positioned according to the extended 10 – 20 system. Two active electrodes attached to the earlobes served as the reference. In addition, we recorded the horizontal and vertical electrooculogram (EOG) using passive electrodes placed at the outer canthi of either eye and above and below the left eye, respectively. EEG was continuously sampled at 512 Hz.

All preprocessing steps were performed using the Fieldtrip toolbox (Oostenveld, Fries, Maris, & Schoffelen, 2011) in the MATLAB environment. First, the EEG data were referenced offline to the average of the earlobe electrodes. Then, data were inspected visually [using the function ft_databrowser()], and bad channels of each participant were rejected. Rejected channels were replaced with a spherical spline interpolation [using the “spline” method in ft_channelrepair()]. Scalp muscle artifacts were detected as epochs in which the amplitude of the bandpass-filtered signal at 110 – 140 Hz at all channels exceeded a z score of 20, with the function ft_artifact_zvalue(). All preprocessing was done on the session as a whole (i.e., without dividing into trials). Division into trials was done for each of the subsequent analyses separately and remaining noisy trials were removed.

Across all trial types, epochs were time-locked to the onset of the tactile stimulus, separately for passive, AE and AO trials. To examine normal stimulus processing, epochs were extracted from 200 ms before to 800 ms after stimulus onset for passive and AE trials. To test for omission trial effects, the same epoch was extracted around an expected stimulus onset (which was not presented) using AO trials. In another analysis, to look for possible pre-stimulus effects, AE trials were epoched from 300 ms before stimulus onset to 800 ms after.

Across all analyses, trial segments were baseline corrected to the mean of the 100 ms period before actual or expected stimulus onset.

### 2.5 Multivariate Pattern Analysis

All MVPAs were performed in Python (V 3.11.3). We used a selection of electrodes over the somatosensory and motor cortex ( ’FT7’, ’FC5’, ’FC3’, ’FC1’, ’FCz’, ’FC2’, ’FC4’, ’FC6’, ’FT8’, ’T7’, ’C5’, ’C3’, ’C1’, ’Cz’, ’C2’, ’C4’, ’C6’, ’T8’, ’TP7’, ’CP5’, ’CP3’, ’CP1’, ’CPz’ ,’CP2’, ’CP4’, ’CP6’, ’TP8’) to train linear discriminant analysis (LDA) classifiers (Grootswagers et al., 2017). This decision was motivated by the fact that the noise artifact from tactile stimulation was strongest in the frontal electrodes and the occipital electrodes are not relevant for the research questions. Each classification analysis was performed at the level of single subjects. It is possible that a similar pattern is elicited by the expectation of touch and actual physical stimulation, but this may occur at different times. Thus, we used temporal generalisation methods whereby a classifier is trained at each timepoint of the training set and tested on each timepoint of the test set. This provides a decoding matrix where each cell represents the average classification accuracy at that train-test timepoint. If the representation of finger location evolves similarly in both train and test sets, above-chance decoding will occur along the diagonal. On the other hand, if the representation unfolds differently over time across train and test sets, above-chance decoding will be off the diagonal.

#### 2.5.1 Within condition decoding

First, we examined whether finger location could be decoded when training and testing within a condition (passive and AE). This involved training an LDA classifier on a portion of within-condition data and testing it on left-out independent trials across all timepoints, using a 5-fold cross validation method. This was done time-locked to stimulus onset (-200 – 800 ms). An above-chance classification performance indicates that the EEG signal contained information that allowed the classifier to discriminate between stimulation to the index or ring finger.

#### 2.5.2 Omission trial decoding

To test for finger specific activity despite the absence of a stimulus, we ran cross-condition decoding analyses. Here, LDA classifiers were trained on passive or AE trials (-200 – 800 ms around stimulus onset) and tested on AO trials (-200 – 800 ms around expected stimulus onset). Cross-validation was not necessary because training and test sets are independent. We also ran an exploratory analysis to probe the effect of condition order on AO trial decoding. We split participants into two groups. The ’Passive First’ group included participants who completed the passive condition *before* the active condition and the ’Passive Last’ group included participants who completed the passive condition *after* the active condition. We then ran the MVPA on each group separately, training on passive trials and testing on AO trials.

#### 2.5.3 Pre-stimulus decoding

To test for the presence of stimulus specific predictive activity, we trained our classifiers on a subset of AE trials including the pre-stimulus and post-stimulus period (-300 – 800 ms) and tested them on the pre-stimulus period of left out AE trials (-300 – 0 ms). Above-chance decoding along the diagonal in the pre-stimulus period indicates that the neural activity before stimulus onset contains information about finger location. Off-diagonal decoding, at post-stimulus training timepoints, indicates that pre-stimulus activity is similar to stimulus-evoked activity.

### 2.6 Statistical inference

We used Bayes factors (BFs) to determine above-chance decoding and at chance decoding (i.e., null hypothesis) at every timepoint within each of the participants using the Bayes Factor R package (Morey & Rouder, 2018) implemented in Python (Teichmann et al., 2021). We set the prior for the null hypothesis at 0.5 (chance decoding) for assessing the decoding results. A half-Cauchy prior was used for the alternative hypothesis with a medium width of r = 0.707. Based on Teichmann et al., (2021), we set the standardized effect sizes expected to occur under the alternative hypothesis in a range between -∞ and ∞ to capture above and below chance decoding with a medium effect size (Morey & Rouder, 2018).

BFs larger than 1 indicate that there is more evidence for the alternative hypothesis than the null hypothesis (Dienes, 2011) with a BF greater than 3 considered as "substantial" evidence for the alternative hypothesis i.e. decoding above- or below-chance level. A BF less than 1/3 is considered substantial evidence for the null hypothesis i.e. chance-level decoding (Jeffreys, 1939; 1961).

### 2.7 Task performance

Participants performed above chance in detecting the tactile stimuli presented at a lower frequency with a mean accuracy of 62 % (SE = 14%) in active trials and 70 % in passive trials (SE = 15%). Reaction times (RTs) were calculated from the time that the low intensity stimulus was delivered until the time that the foot pedal was pressed. The mean RT was 714 ms (SE = 132 ms) in the passive block and 754 ms in the active block (SE = 43 ms).

## 3 Results

### 3.1 Within condition decoding of stimulation location (index vs ring finger)

We found that the classifier was able to accurately determine which finger was stimulated at a negligible latency after stimulus onset in both passive (Fig. 2a) and AE (Fig. 2b) trials. The initial decoding is likely too short for neuronal conduction delays, suggesting that very early above-chance decoding may be, at least partially, informed by an artifact of the mechano-vibratory stimulus. This is further elaborated on in the discussion section. Despite this, above-chance decoding was sustained well beyond the stimulus duration. In the passive condition, there was sustained decoding of finger location lasting until 800 ms after stimulus onset. In the expected condition, there is strong evidence for finger location decoding up until ∼400 ms after stimulus onset. Following that point, decoding is slightly less consistent, although there is evidence of a finger location representation up until 800 ms. These results clearly demonstrate that multivariate classifiers were able to extract finger location information from the EEG signal.

**Figure 2.**
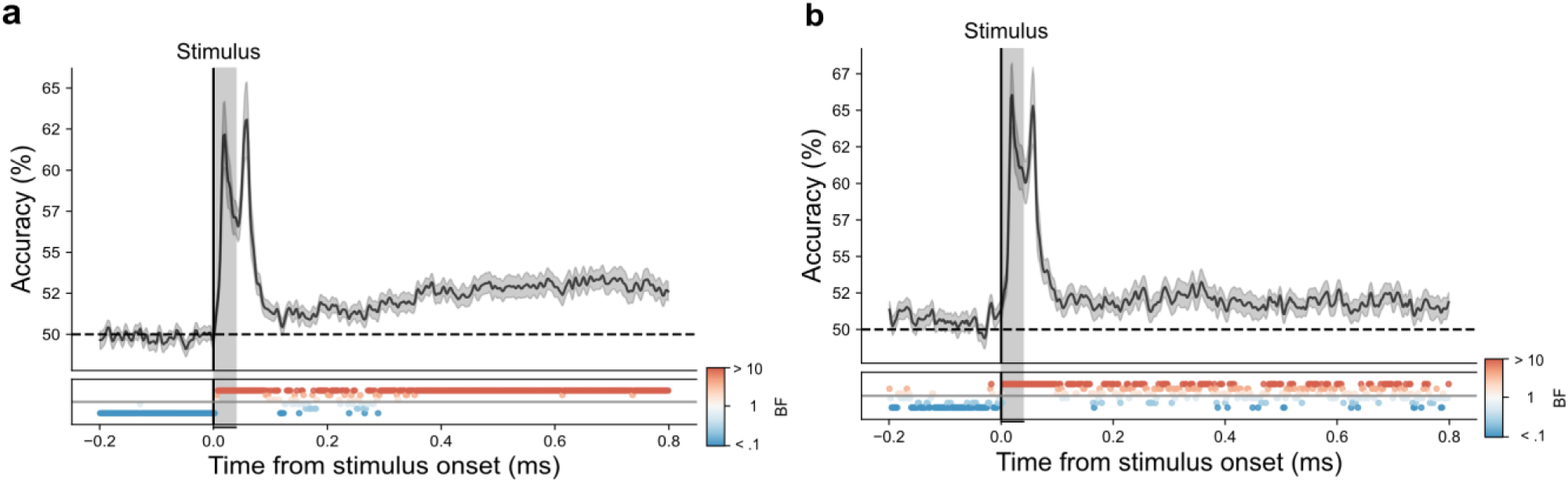
**Above-chance decoding of finger location in passive and expected trials** Classifiers were trained and tested to distinguish patterns of neural activity evoked by stimulation of different fingers for **a** passive trials and **b** AE trials. The plotted results reflect the mean participant classification performance N = 35) at corresponding training and testing timepoints; the shaded area represents the standard error. The grey vertical section shows the duration of the vibration stimulus (40 ms). The Bayes factors (BF) below the plots indicate the timepoints at which there was substantial evidence in favour of the alternative hypothesis i.e. decoding above-chance (dark red) and substantial evidence for the null hypothesis, i.e. chance level (dark blue).

### 3.2 Stimulus omissions evoke a finger-specific prediction error

When training on AE trials and testing on AO trials, there was strong evidence that information about expected stimulation location was present in the neural activation patterns.

This occurred between ∼120 and ∼250 ms following anticipated stimulus onset time, despite the absence of a physical stimulus (Fig. 3). This finding indicates that neural activation patterns evoked by the absence of expected finger stimulation is similar to stimulus-evoked activity. Importantly, this overlap occurred primarily along the diagonal of the temporal generalization matrix. This means that signals associated with the predicted stimulus unfolded at a similar time course as the actual processing of the stimulus.

**Figure 3.**
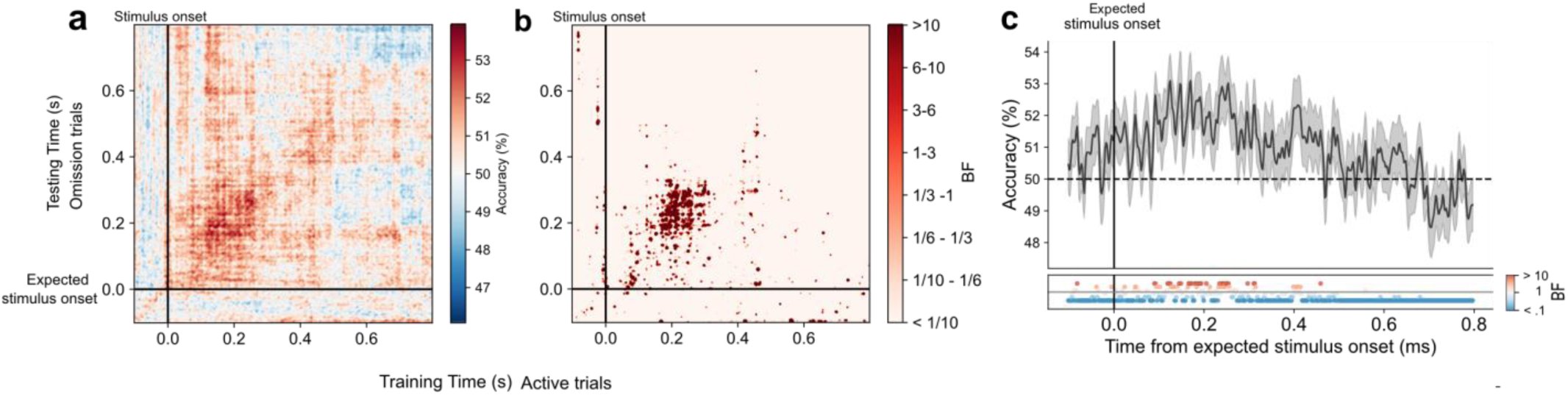
**Finger-specific prediction error on AO trials (training on AE trials)** Classifiers were trained on AE trials to distinguish patterns of neural activity evoked by stimulation of different fingers and subsequently tested on AO trials. **a.** Temporal generalisation matrix (TGM) showing above-chance decoding of finger location after omission onset. **b.** The corresponding bayes factors. BFs show the strongest evidence (red) when training and testing on corresponding timepoints i.e., along the diagonal. **c.** Decoding along the diagonal of the TGM shown in a. The group average (N = 35) decoding accuracy is plotted as a function of time from expected stimulus onset in omission trials; the shaded area represents the standard error. The Bayes factors (BF) indicate the timepoints at which there was substantial evidence in favour of the alternative hypothesis i.e., decoding above-chance (dark red) and substantial evidence for the null hypothesis, i.e. chance level (dark blue).

In AE trials participants had full control over stimulus onset time, making stimuli predictable in both time and location. Consequently, while the training data included stimulus-evoked activity, it likely also contained predictive signals since participants could anticipate the upcoming stimulus. This complicates the conclusions we can draw about the omission representation as it is not clear whether the trained classifier is relying on bottom-up signals generated by stimulus presentation or top-down activity arising from stimulus anticipation.

To remove the influence of predictive signals and isolate the contribution of bottom-up stimulus activity, we trained our classifiers on passive trials in which the stimulus was unpredictable in both timing and location. Trained classifiers were tested on AO trials. We found almost exclusively at-chance decoding as shown in Fig 4. In other words, expected finger location could not be extracted from the omission response when training on data from passive trials. This may indicate that above-chance decoding shown in Fig. 3 is driven by prediction-related activity.

**Figure 4.**
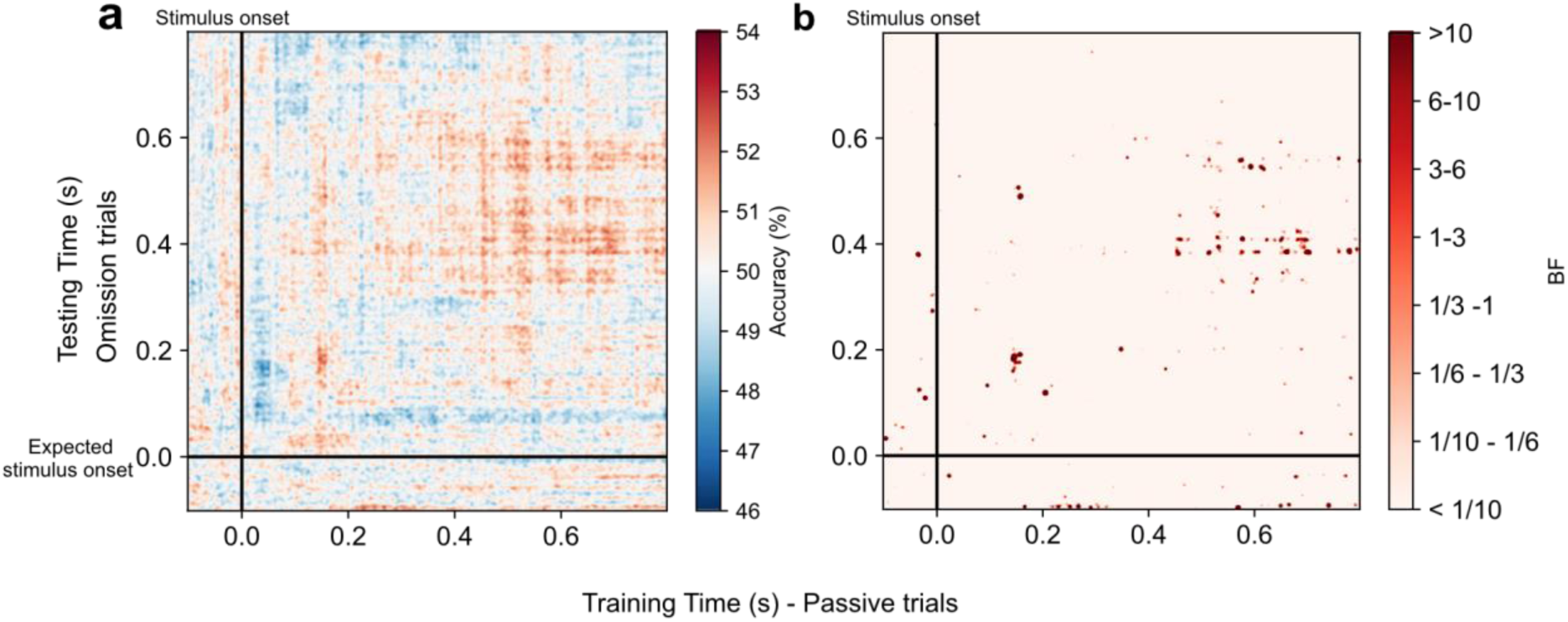
**The omission response cannot be decoded when training on passive trials** Classifiers were trained on passive trials to distinguish patterns of neural activity evoked by stimulation to the index versus the ring finger and subsequently tested on AO trials. **a.** Temporal generalisation matrix (TGM) showing above-chance decoding (> 50%) of finger location after omission onset. **b.** The corresponding Bayes factors. BFs show very little evidence for decoding of finger location, with mostly strong evidence for chance decoding (white).

### 3.3 Exploratory analysis of omission response (training on passive)

In an exploratory analysis, we tested for the effect of block order when training on passive trials and testing on AO trials. Decoding results revealed above-chance decoding of finger location only for the ‘Passive Last’ group. Strong evidence for above-chance decoding of finger location emerged ∼100 ms after omission onset at training timepoints between ∼200 and ∼800 ms (Fig, 5). This indicates that the omission response was similar to a late representation of the physical stimulus as presented on passive trials.

### 3.4 Action induces predictive tactile stimulus templates

Two interesting findings emerged when training on a subset of AE trials and testing on the pre-stimulus period of a separate subset of AE trials. Firstly, we found above-chance decoding along corresponding training and testing timepoints in the pre-stimulus period (Fig 6a. data within black bands) and secondly, we found pre-stimulus decoding when using late post-stimulus training data (Fig 6a. data within green bands). For each participant, we calculated the average classification score within the bands. For the diagonal band in the pre- stimulus period there was strong evidence for above-chance decoding of finger location from 300 – 100 ms before stimulus onset (Fig 6b. *top panel)*. This means that there was information about the expected stimulation location in pre-stimulus EEG activity. In addition, we calculated classification performance in the pre-stimulus period averaging over training times from 300 – 800 ms post stimulus onset. The strongest evidence for above-chance classification performance was between ∼150-130 ms before stimulus onset (Fig 6b. *bottom panel*). This suggests that starting at ∼150 ms the predictive representation of finger location becomes similar to the late representation of a physically presented tactile stimulus.

**Figure 5.**
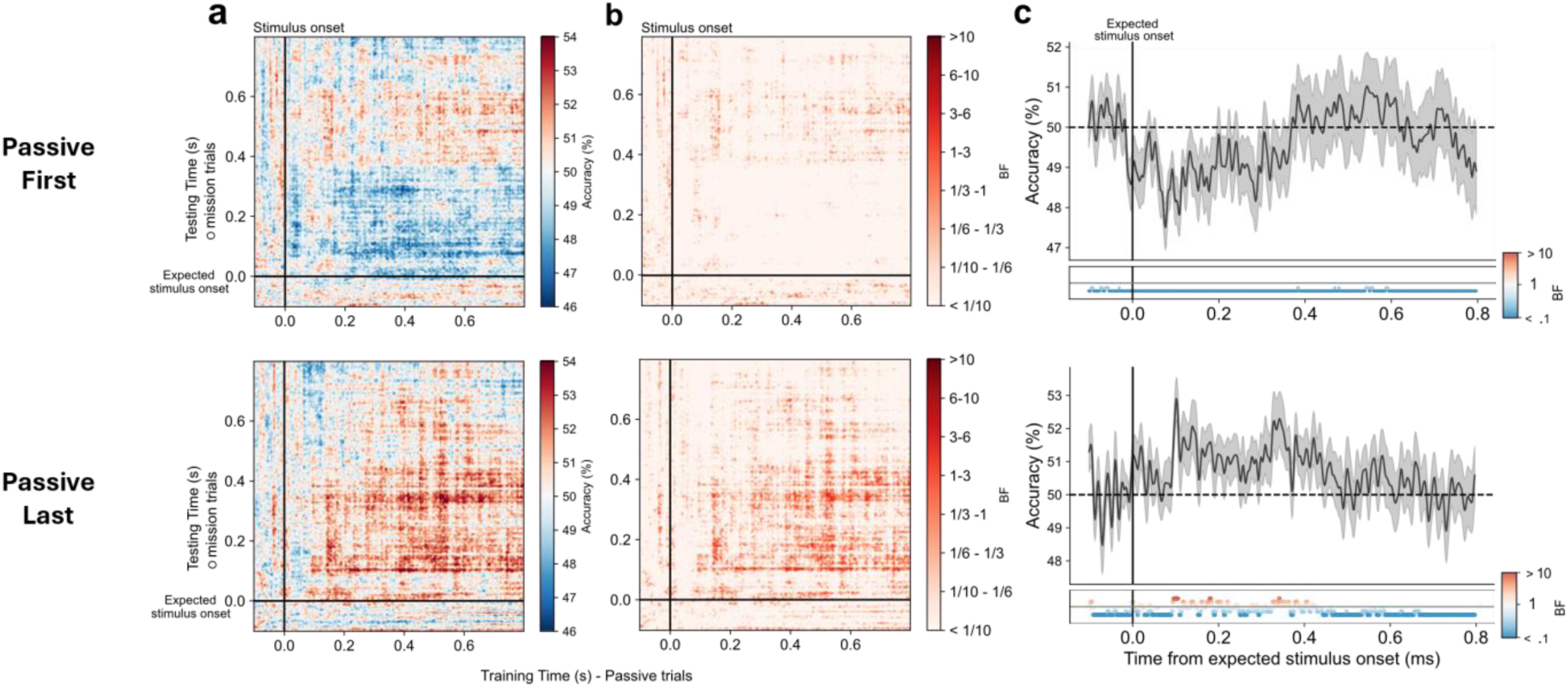
**Block order affects decoding of omission response (training on passive trials)** Classifiers were trained on passive trials to distinguish patterns of neural activity evoked by stimulation of different fingers and subsequently tested on AO trials. Results are grouped into participants who did the passive block before the active block (Passive First – Top panel) and participants who did it after (Passive Last -Bottom Panel) **a.** Temporal generalisation matrices (TGMs) showing above-chance decoding of finger location after omission onset for Passive First (top panel) and Passive Last (bottom panel) groups. **b.** The corresponding Bayes factors. BFs show mostly strong evidence for chance decoding (white) in the Passive First group while the Passive Last group show evidence for above-chance decoding (red) between 100 and 400 ms (using training timepoints between 200 and 800 ms). **c.** The group mean (N = 35) decoding accuracy is plotted as a function of time from expected stimulus onset in omission trials averaged across training timepoints 200-800 ms after stimulus onset of passive trials. The shaded area represents the standard error. The Bayes factors (BF) indicate the timepoints at which there was substantial evidence in favour of the alternative hypothesis i.e., decoding above-chance (dark red) and substantial evidence for the null hypothesis, i.e. chance level (dark blue).

**Figure 6.**
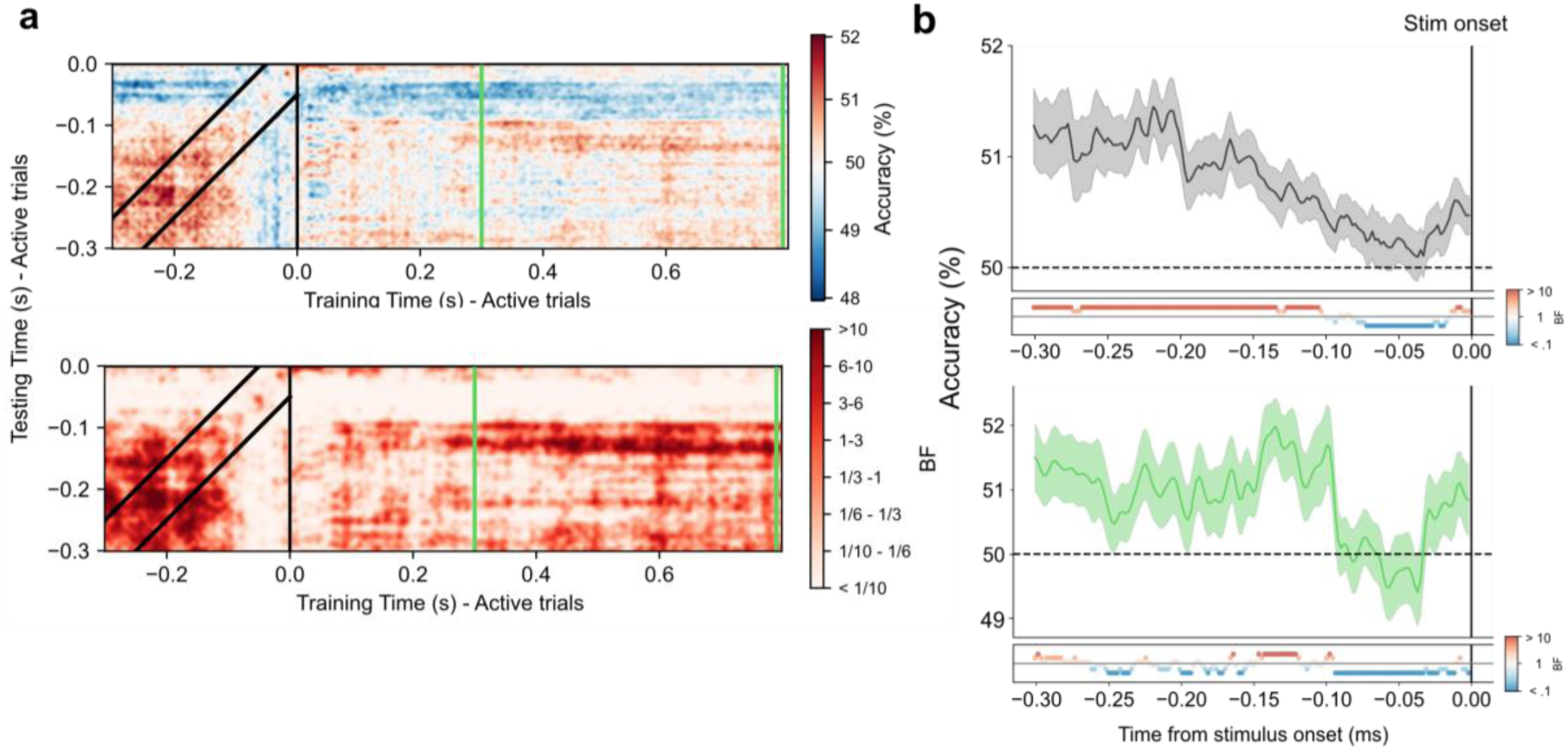
**Finger-specific tactile template in the pre-stimulus period (training on expected trials)** **a.** *Top panel* Temporal generalisation matrix (TGM) of classifiers trained on the time course of AE trials and tested on the pre-stimulus period of left-out AE trials. ***Bottom panel***: The corresponding Bayes Factors for the TGM. BFs show the strongest evidence (red) when training and testing along the diagonal of the pre-stimulus period (illustrated by the black diagonal bands) and at training times from 300 to 800 ms after stimulus onset (illustrated by the vertical green bands). **b.** For each participant, we calculated the average classification performance in two bands (shown in a). The black diagonal band reflects the similarity between corresponding training and testing times in the pre-stimulus period (***top panel***). The green band reflects the similarity between stimulus-evoked activity from 300-800 ms and pre-stimulus activity (***bottom panel***). The group mean (N = 35) decoding accuracy is plotted as a function of time before stimulus onset; the shaded area represents the standard error. The Bayes factors (BF) below the plots indicate the timepoints at which there was substantial evidence in favour of the alternative hypothesis i.e., decoding above-chance (dark red) and substantial evidence for the null hypothesis, i.e., chance level (dark blue).

## 4 Discussion

As embodied agents, we can manipulate the environment through action to produce desired effects. This is true for effects distal from the body such as typing on a keyboard to produce a sentence, as well as more proximal, body-related effects such as scratching an itch i.e., self-touch. In both cases, an anticipatory representation of the upcoming sensory consequences is formed, i.e., words appearing on the screen or the tactile sensation of the scratch. This acts as a template against which feedforward sensory information is compared. When the sensory input differs from what was predicted, a prediction error is elicited. This includes when the expected action-effect is absent entirely, for example, if the keyboard is not plugged in.

It has been documented that the somatosensory cortex can be activated even without afferent input; for example, during touch observation or tactile imagery (Kuehn et al., 2014; De Borst & De Gelder, 2017) and a recent study demonstrated pre-stimulus somatotopic activity in anticipation of a vibrotactile stimulus (Kassraian et al., 2023.). It is however, not known whether motor-induced expectation of a tactile stimulus elicits similar somatotopic activity. To investigate action-induced tactile predictions, we asked participants to make index finger movements which were paired with stimulation to the index or ring finger (block dependent) of the opposite hand. On rare trials, the tactile stimulation was omitted entirely. Thus, within each block participants formed a location-specific tactile prediction, with stimulus timing fully controlled by participants’ movements. Recording EEG and applying a multivariate analysis approach to the data we found that in the absence of bottom up input we could decode expected finger stimulation location, effectively capturing a location-specific prediction-error response. In addition, we found that pre-stimulus activity contained information about expected stimulation location, evidence of an anticipatory tactile template. This is, to the best of our knowledge, the first study to show neural evidence of finger-specific tactile predictions, driven by self-generated actions.

### 4.1 The omission response

We were interested in the presence of finger location information in neural activity following an omission. Given that omission-related responses cannot be explained by feedforward activity along the ascending somatosensory pathway, information about stimulus identity is likely attributable to a purely prediction-related response (Wacongne et al., 2012). Our results illustrate that, following a stimulus omission, finger-specific information is present approximately 120 - 250 ms later. This is in line with previous reports from the auditory domain which find an omission induced ERP, emerging between 100 – 150 ms (Todorovic, 2011; Wacongne et al., 2011; SanMiguel, Widmann et al., 2013; Sanmiguel et al., 2013).

The omission response can be explained by predictive coding whereby the repeated coupling of a finger movement with the finger-specific tactile stimulus allows a sensory prediction to be formed. This prediction is sent (perhaps as a corollary discharge signal) through descending connections to sensory areas, activating neurons tuned towards the expected stimulation location (i.e. index or ring finger) (Reznik et al., 2015; Schneider & Mooney, 2018; Pazen et al., 2020).

While above-chance decoding of finger location on omission trials strongly supports the presence of stimulus specific predictions, distinguishing whether this activity reflects the prediction itself or a prediction error signal remains challenging. Predictive coding models assume that, on omission trials, the prediction (e.g. vibration to the index finger) is subtracted from the bottom up input (no vibration), producing a negative prediction error. Negative prediction errors arise when the actual input is weaker than predicted, such as during an omission, and may be neurally implemented through bottom-up inhibition of the relevant neurons (Keller & Mrsic-Flogel, 2018). This inhibition likely suppresses activity of neurons encoding the expected stimulus. Despite this, we report above-chance decoding of finger location. This indicates that omission decoding may reflect the sensory prediction of the absent stimulus or perhaps a combination of prediction-related activity and error-related activity (Wacongne, 2012; Walsh, 2020). Thus while we can conclude that tactile representations are predictively evoked by movement, further research is needed to identify the precise neural origins of the omission response.

Under the predictive coding framework, prediction-error signals are thought to be stimulus-specific and not simply a ‘surprise’ signal, (Kok & De Lange, 2015). This necessitates a precise representation of upcoming input. MVPA has previously been used to determine the representational contents of prediction signals, although outside the action domain and most often within the visual and auditory modalities (Kok et al., 2012, 2017; Demarchi et al., 2019). In an influential study, fMRI was recorded while participants viewed oriented gratings. The orientation of the grating was fully predictable by the preceding tone, either at a low- or high-frequency (Kok, 2014). On trials in which the grating was unexpectedly omitted, the expected orientation could be decoded from V1 activity. Later work using MEG demonstrated that predictive signals about the upcoming stimulus were already present 40 ms before its presentation (Kok et al., 2017) supporting the existence of predictive stimulus-like templates induced by expectation.

The fact that omission-related activity could be decoded by classifiers trained on physically presented stimuli indicates that the neural representation of the expected stimulus resembled the neural activity evoked by the physical stimulus. Across two different analyses, we used AE trials (active trials with a stimulus) and passive trials to train our classifiers; testing them on AO trials (active trials without a stimulus) with somewhat different results. The first interesting difference is the training times at which above-chance decoding emerged when training on AE versus passive trials (‘Passive Last’). For AE trials, above-chance omission decoding was found along the diagonal meaning that at corresponding training and testing timepoints the activation pattern is similar. In contrast, when training on passive trials, late tactile patterns were the most similar to activity elicited by the omission. More precisely, the omission response contained information that occurred ∼400 ms after being stimulated in passive trials. The difference between training on passive versus AE trials may be explained by the inherent predictability in AE trials. In passive trials, the tactile stimulus was unpredictable in terms of both finger location and onset time. This is in contrast to AE trials where location and timing were fully predictable. The formation of a prediction in both training (AE) and test sets (AO) may explain the similarity of representations at corresponding timepoints.

When training on passive trials we found that the location-specific omission response could only be decoded in the ‘Passive Last’ group. This phenomenon may be explained by ideomotor theory. The ideomotor theory proposes that repeated exposure to actions and their corresponding sensory effects leads to a bidirectional relationship such that both action and its consequences are represented by a common code (Eisner & Hommel, 2001; Hommel, 2013; Shin et al., 2010). Thus, activating the representation of an action-effect automatically triggers the action itself. Our finding supports the idea of a common code whereby after repeated exposure to the index finger movement and the corresponding tactile stimulation, presentation of the stimulus alone elicits a pattern of activation that overlaps with the movement itself. This is an interesting finding under the current conditions as the movement was equally paired with stimulation to the index and ring finger. This suggests that multiple action-effect pairing can be held simultaneously and flexibly employed, perhaps adapting as the context changes.

### 4.2 The predictive stimulus template

Our results support the idea that prediction-related top-down activity pre-activates stimulus-specific neural ensembles (Ekman et al., 2017; Kok et al., 2017), further showing that this is the case for movement-related predictions. We found that when training and testing on the pre-stimulus period (AE trials), a sustained representation of upcoming finger location was present from 300 to 100 ms before stimulus onset. Crucially, this above-chance decoding cannot be attributed to the movement itself, as the movement was identical for both index and ring finger stimulations. From ∼150-100 ms before stimulus onset the predictive representation becomes more stimulus-like, evidenced by above-chance decoding using training timepoints from 300 – 800 ms post-stimulus. This is similar to the predictive templates reported in the visual domain, whereby expected orientation could be decoded from MEG before its onset (Kok et al. 2017). In contrast to Kok’s study we find evidence for a predictive template long before the stimulus and this disappears 100 ms before its onset, likely around the start of movement preparation and execution.

In our analysis we epoched around the onset of the stimulus instead of movement onset to provide more accurate time-locking. That said, it is highly likely that the anticipatory representation of finger location is available long before movement onset. Movement onset was determined by a distance threshold whereby when participants’ index finger descended 10 mm, we considered it the start of the movement. We chose a liberal threshold as we did not want small, unintentional movements to trigger the onset of the vibration. That said the average time between the finger passing 10 mm and the stimulus being delivered (5 mm later) was 30 ms. The velocity profile of voluntary movements is usually bell-shaped, symmetric and contains a single peak (Morasso 1981; Abend et al. 1982; Atkeson & Hollerbach, 1985; Flash & Hogan, 1985; Uno et al., 1989). In other words, there is an initial acceleration phase before reaching peak velocity which is then followed by a deceleration phase. We know that participants took 30 ms to move their finger 5 mm, likely in the deceleration phase. However even if we assume the finger moved at a constant speed then the total time from initiation to stimulus would be 90 ms. This is probably an overestimation given acceleration and deceleration times. There is no evidence of above-chance decoding in the 100 ms period before stimulus onset, likely when the finger is in motion. In fact, we find a steep drop in decoding accuracy 100 ms before stimulus onset. Thus, these results support the presence of a predictive stimulus-like template *before* movement onset.

### 4.3 Non-motor prediction mechanisms

In the literature, action-effect predictions are most often described under a forward model framework (Wolpert & Miall, 1996). While self-generated touch is no exception (Blakemore et al., 1998), our findings violate some of the tenets associated with a forward model. One major assumption of the forward model is that updating internal predictions occurs gradually (Rummell et al., 2016; Schneider et al., 2018). In the current experimental setup, the action-effect pairing alternated from block to block. This means that in one block finger movement caused a vibration to the index finger while in the next block the vibration came to the ring finger. Despite this, we find reliable above-chance decoding of finger location in the pre-stimulus period and on omission trials. This demonstrates a rapid and flexible reconfiguration of predictions whereby the brain updates the action-effect pairing depending on the context. The gradual updating described in previous animal studies, where attenuation for expected action-outcomes only emerged after extensive practice over several days (Rummell et al., 2016; Schneider et al., 2018), contrasts sharply with our findings. This difference indicates that motor-based forward models may not fully account for the rapid prediction updating we observed.

Another important attribute of the forward model is that prediction depends on how likely it is that the sensation was self-generated. Previous behavioural studies have shown that the attenuation of self-touch is determined by the causal relationship between the movement of the finger and the resulting sensation (Bays & Wolpert, 2008; Kilteni & Ehrsson, 2017). For instance, when hands are positioned such that self-touch is impossible, the sensory attenuation effect is significantly reduced (Bays & Wolpert, 2008) or absent entirely (Kilteni & Ehrsson, 2017). This indicates that tactile attenuation is modulated by the plausibility of the sensation being self-generated. When preparing a movement, the forward model, using the efference copy and the current position of the hand, predicts the future position of the moving index finger. If the future position does not align with the passive index finger to ultimately allow contact, the model does not anticipate contact between them, and thus no attenuation ensues (Kilteni & Ehrsson, 2017). In our setup, while the moving and passive hands were relatively close to one another, the distance did not allow for real self-touch.

Nevertheless, we found strong evidence for prediction formation about stimulation location. These observations suggest that the forward model may be more flexible than originally conceived. Another possibility is that our results can be explained by non-motor mechanism.

Related research demonstrates that sensory events can be predicted using statistical regularities in the environment. This is demonstrated through attenuation effects for non-motor predictions across the visual (Alink et al., 2010; Kok et al., 2012; Summerfield & Koechlin, 2008; Richter et al. 2018) and auditory (Lange, 2009; Hughes et al., 2013; Schröger et al., 2015) modality. This supports the idea that sensory input can be predicted using stimulus probabilities (de Lange et al., 2018). Some have suggested that the motor system may be part of a more general prediction system that guides both perception and action (Thomas et al., 2022; Press et al., 2023). This is in line with theoretical accounts such as active inference (Friston et al., 2009; Friston et al., 2011; Friston, 2011; Adams et al., 2013 and the ideomotor theory (Eisner et al., 2001; Shin et al., 2010; Hommel, 2013). While these also posit that prediction of action consequences accompanies an action, they do not rely on a forward model. It is possible that the current findings can be explained by a more general prediction mechanism, independent of motor-specific forward models. The relationship between motor- and non-motor predictions is unclear. As is expected, non-motor predictions are usually assessed in contexts that do not involve action, making direct comparisons challenging. Our study manipulated prediction within the context of self-generated action taking steps to link ideas from outside and inside the action domain (Thomas et al., 2022).

Future work should investigate how non-motor predictions influence motor-perception interactions and establish whether the same neural mechanisms generate predictive stimulus templates inside and outside the action domain.

## 5 Conclusion

In this study, we investigated the prediction of self-produced tactile events, providing novel evidence for finger-specific predictions in the somatosensory domain. We found that when a tactile stimulus was expected but omitted, the omission response contained finger-specific information. This supports predictive coding theories where the prediction error constitutes the mismatch between the prediction and actual sensory input. In addition, we found evidence for predictive activity in the pre-movement period, with a sustained representation of expected finger location. As movement approached, pre-stimulus EEG activity became more stimulus-like, mirroring findings from non-action related research. This indicates that the brain generates anticipatory stimulus templates against which feedforward information can be compared, also in action contexts. By employing methods comparable to those used in studies outside the action domain, our research highlights similarites between action-induced and environment-induced predictions. These insights advance our understanding of the predictive mechanims that support action, bringing us closer to comprehending how the brain prepares us to optimally engage with the environment.

## Acknowledgments

We thank the participants of the study, Clare Press and Emily Thomas for providing experimental code related to programming vibrotactile stimuli, Mauro Ejzenberg for technical support with the vibrotactile equipment, members of the Landau Lab for participating in early pilots and Talya Shlezinger for collecting the data.

